# Mechanisms of Social Behavior in the Anti-Social Blind Cavefish (*Astyanax mexicanus*)

**DOI:** 10.1101/2024.11.30.626194

**Authors:** Britney Sekulovski, Noam Miller

## Abstract

The evolution of social behavior in *Astyanax mexicanus*, which exists as a sighted, surface-dwelling morph and a blind, cave-dwelling morph, provides a model for understanding how environmental pressures shape social behaviors. We compared the shoaling behavior of blind and surface *A. mexicanus* to that of zebrafish (*Danio rerio*), and examined the effects of nutritional state and the neuropeptides isotocin (IT) and arginine vasotocin (AVT) on their social behavior. Blind cavefish not only fail to form shoals, but actively avoid conspecifics, with hunger further diminishing their social cohesion. Administration of low doses of AVT and an IT antagonist partially restored social behavior in blind cavefish, reducing distances between individuals, whereas surface fish exhibited minimal or opposite responses to these hormonal manipulations. Our findings suggest that the loss of schooling behavior in blind cavefish is not a consequence of visual impairment alone, as they remain capable of detecting and responding to others. Instead, this behavior likely reflects an adaptive response to their resource-poor, predator-free cave environment, where shoaling may be disadvantageous. The differing responses to social hormones between the morphs indicate that blind cavefish may have lost the motivation to shoal rather than the ability, highlighting how ecological pressures can shape social behavior.

## Introduction

Social behaviors vary widely across species, often shaped by diverse environmental pressures. Among vertebrates, fish exhibit a particularly extensive range of social behaviors, from predominantly solitary species (1) to those that engage in cooperative breeding (2), establish dominance hierarchies (3), or even display reciprocal altruism (4). Most social fish species engage in collective movement, a behavior often referred to as shoaling (5–8). Shoaling offers significant fitness benefits, including predator avoidance, as it increases vigilance, enabling quicker predator detection and creates confusion for predators when faced with multiple identical targets (9–16). Shoaling also enhances foraging efficiency by enabling fish to locate and exploit larger food patches and increasing feeding efficiency due to reducing the need for individual vigilance (15,17–23). However, aggregation can also have costs, such as increased competition for limited resources (24,25) or an elevated risk of attracting predators to large groups (12,13,26).

The balance between the benefits and costs of shoaling raises intriguing questions about the evolution of this behavior and how environmental conditions shape it (27). For instance, the presence of predators has been shown to impact the evolution of social and anti-predatory behaviors (28). Trinidadian guppies (*Poecilia reticulata*) derived from populations originating in low-predator environments display reduced social cohesion compared to descendants of fish from high-predation areas (29–35). Similarly, desert pupfish (*Cyprinodon macularius*), which evolved in predator-free environments, appear to have lost defensive behaviors such as responses to alarm pheromones (36).

Variations in the benefits of aggregation can also lead to adjustments in collective behaviors in response to current environmental conditions. Factors like pollution, waterway obstruction, and noise modify group cohesion (37–40). Zebrafish (*Danio rerio*) exposed to simulated predator risks form denser schools (5). Gulf menhaden (*Brevoortia patronus*) form tighter, more synchronized groups in physically complex environments than in open water (41), while zebrafish exhibit the opposite pattern (42). Nutritional state further influences social cohesion, with hungry zebrafish preferring to shoal with well-fed conspecifics, potentially to increase their chances of finding food while minimizing competition (43). When food is present, zebrafish also increase their distance from neighbors (5). Food deprivation reduces shoaling tendencies across species; food-deprived banded killifish (*Fundulus diaphanus*), walleye pollock (*Theragra chalcogramma*) and three-spined sticklebacks (*Gasterosteus aculeatus*) display a reduced tendency to maintain tight shoals and display less cohesion with conspecifics (44–46). Similarly, European sea bass (*Dicentrarchus labrax*) exhibit fewer interactions with others when food-deprived (47).

The Mexican tetra (*Astyanax mexicanus*, AM) is a valuable model for studying the evolution of social behavior in fish. Approximately 20,000 years ago, some surface-dwelling populations of this species migrated into pitch-black caves, where they evolved into a blind cave-dwelling form (48–51). This transition occurred multiple times across up to 34 different caves (52), with populations independently evolving the same traits, providing a striking example of convergent evolution (53–56). Blind AM populations underwent a host of morphological, physiological, and behavioral adaptations (57–66). These changes are believed to have been driven not only by the complete absence of light but also by the lack of predators and extreme scarcity of food in their cave habitats. In such habitats, blind AM feed on low-nutrition organic matter that occasionally drifts into the caves, such as detritus, algae, fungi, bat guano, and the remains of other cave-dwelling organisms (51,53,67–70). Many populations of blind AM, such as Pachón cave populations, are characterized by their relentless pursuit of food and have been suggested to be insatiable (57,60).

In addition to losing their eyes and pigmentation, cave-dwelling blind AM have also evolved an enhanced lateral line system, a sensory adaptation crucial for survival in the dark (54,71–73). The lateral line, an array of pressure-sensitive cells along the sides of fish, acts as a sensory organ that detects the velocity of water flow generated by the fish’s own movements or by external currents (74–76). Blind AM possess a higher density of these cells, which improves their sensitivity to water movements, aiding in the detection of food and obstacles (77–84). They also use active sensing, generating a stable flow as they swim, allowing them to detect distortions in the water caused by nearby objects (85–87).

Although born with eyes, blind AM undergo lens apoptosis early in development, a process driven by increased expression of the *sonic hedgehog* (*Shh*) gene (88–90). Elevated *Shh*, alongside Fibroblast Growth Factor (*Fgf*) signaling, contributes to many of the adaptations seen in blind AM, such as larger jaws, an increased number of taste buds, and expanded forebrain regions, including the olfactory system and hypothalamus, which result in a higher number of neurons associated with feeding behavior (91–94). Consequently, these neuroanatomical changes cause blind AM to sleep less, spend more time foraging, and have heightened appetites— adaptations that enhance their ability to maximize food intake and forage more efficiently than surface fish in the dark (57,59,60,70,90,95).

Alongside these physiological adaptations, blind AM have undergone several behavioral changes (72). The most notable behavioral shift is their complete loss of shoaling behaviors (53,60,64,67,69,96–100). Although this change has often been attributed to their loss of vision (53,64,96), it is essential to consider the broader ecological context: in addition to scarce food, blind cavefish also have no predators (53,67). Therefore, the adaptive benefits of shoaling, such as predator avoidance and collective foraging, are greatly reduced in their environments. Here, we propose that the loss of shoaling in blind AM results more from a decrease in their motivation to shoal than an inability to aggregate.

While shoaling remains a vital survival mechanism for surface-dwelling AM, providing protection from predators and enhancing foraging efficiency (50,53,96), solitary foraging might be a more effective strategy in habitats with high competition for limited resources (101,102). Swimming independently might also improve lateral line function in blind AM by enhancing their signal-to-noise ratio, aiding navigation and foraging in complete darkness (60,81). Blind AM also exhibit a reduced response to alarm substances, further suggesting that the absence of predators has relaxed the selective pressures maintaining shoaling and defensive behaviors in their surface-dwelling relatives (64,103). Even hybrids of surface and cave morphs exhibit reduced shoaling, despite retaining functional vision, indicating that inherited predispositions from blind AM, rather than vision loss alone, contribute to this behavioral shift (64).

Alongside the loss of shoaling, blind AM show a significant reduction in aggressive behaviors, such as decreased biting and a complete loss of territoriality and hierarchical dominance. In contrast, surface AM exhibit hierarchical dominance and high levels of aggression (60,69,96–98,104). As with shoaling, this loss of agonistic behaviors in blind AM was initially attributed to a loss of vision, as aggression was thought to rely on visual cues (96,97,105). However, subsequent studies have demonstrated that surface-dwelling sighted AM remain aggressive even in complete darkness (60), and hybrid populations also exhibit reduced aggression (106). This behavioral shift likely stems from changes in food-seeking behavior and neurobiological adaptations, such as alterations to the serotonergic system (60,98,104).

Hormones likely play a crucial role in shaping social behaviors in blind AM, as they do across many fish species (107,108). In blind AM, adaptations in hypothalamic brain regions driven by *Shh* and *Fgf* signaling not only enhance their foraging abilities but also contribute to changes in their neuroendocrine system by reducing arginine vasopressin (AVT)-producing neurons (94). AVT, known for its role in stress regulation (109), may contribute to the distinct behaviors of blind AM, such as their lower baseline cortisol levels, compared to surface AM in familiar environments (110). As AVT also influences social behavior, these neuroendocrine changes may help explain blind AM’s reduced shoaling and decreased motivation for group living. Examining social hormones like AVT could thus offer valuable insights into the mechanisms underlying social behavior—or its absence—in this species.

Neuropeptides such as AVT and isotocin (IT)—the fish analogs of the mammalian arginine vasopressin (AVP) and oxytocin (OT) —play essential roles in regulating social behaviors across vertebrate taxa (111,112). In goldfish (*Carassius auratus*), IT decreases proximity to conspecifics, while AVT promotes shoaling (113). In guppies, IT has the opposite effect: it increases social proximity, while AVT decreases grouping (114). However, another study on guppies found that IT increased social cohesion, while AVT was linked to anxiety-like behaviors without affecting shoaling (115). In zebrafish, AVT reduces social interactions and aggression (116,117), but it increases aggression in species such as bluehead wrasse (*Thalassoma bifasciatum*; 118,119), brown ghost knifefish (*Apteronotus leptorhynchus*; 120), and beaugregory damselfish (*Stegastes leucostictus*; 121), highlighting species-specific differences in neuropeptide effects.

Cichlids also show varied responses: in male *Neolamprologus pulcher*, IT reduces grouping, yet increases sensitivity to social stimuli, while an IT receptor antagonist promotes grouping (122,123). *N. pulcher* also has higher brain expression of IT-related genes linked to social behaviors than the non-social cichlid *Telmatochromis temporalis* (124). In African cichlids (*Astatotilapia burtoni*), AVT helps maintain social hierarchies, with dominant males showing higher AVT expression in key brain areas than subordinates (125,126). Environmental context also influences AVT’s role in social dynamics. For instance, guppies exposed to predation risk show increased AVT expression without a corresponding change in IT expression, suggesting that predator presence may specifically enhance AVT’s role in social regulation (127). In mosquito fish (*Gambusia affinis*), IT modulates social behaviors in context-specific ways, such as reducing interactions with males while maintaining associations with conspecific females under conditions of male harassment (128). This supports the social salience hypothesis that OT/IT increases the salience of social stimuli—both positive and negative—thus adapting social behavior to environmental context (128–130).

In mammals, OT enhances positive social interactions by increasing proximity in lions (131), partner-seeking behavior in marmosets (132), and caregiver attention in infant macaques (133). In vampire bats, OT increases food donation size and allogrooming (134), in naked mole-rats, it boosts huddling and proximity to familiar conspecifics (135), and in meerkats, it increases cooperative behaviors, such as pup-feeding, while reducing aggression (136). In humans, OT is linked to trust, empathy, and social connectedness (137–139). In contrast, AVP is more often linked to behaviors related to withdrawal, such as aggression, territoriality, and defense. For instance, AVP increases aggression in male rodents, bolstering their territorial and mate-defense behaviors (140). In avian species, AVP analogs help maintain social bonds and manage dominance hierarchies, similar to their roles in fish (141). Despite the substantial influence of AVP/AVT and OT/IT on diverse social behaviors, their role in complex social dynamics, particularly in shoaling behaviors, remains poorly understood.

To investigate whether the loss of shoaling in blind AM represents an adaptive strategy rather than a physiological constraint, we examined the shoaling tendencies of surface-dwelling and cave-dwelling AM morphs alongside zebrafish—a well-studied schooling species used as a control. We also compared these groups to a theoretical shoaling-null model that assumed they were swimming randomly and ignoring one another. We hypothesized that blind AM would exhibit reduced shoaling behaviors compared to sighted AM and zebrafish, reflecting an adaptive response to their resource-scarce, predator-free cave environments. Next, we manipulated the nutritional state of blind AM to examine the effects of hunger on shoaling. Fish were observed in three states: fasted (24 hours after feeding), post-absorptive (approximately 3-5 hours after feeding), and fed (10 min after feeding). We hypothesized that hunger would reduce social cohesion while feeding would enhance it, as it does in other species (44–47). Finally, we administered varying doses of IT and AVT, as well as their antagonists, to both AM morphs, to explore the role of these neuropeptides in modulating shoaling behavior. We hypothesized that the hormonal responses would differ between morphs, reflecting their contrasting reliance on social interactions and differences in their neuroendocrine regulation. Together, these experiments aim to reveal how adaptations to extreme ecological conditions, like the complete absence of light, contributed to the loss of shoaling in cave-dwelling populations, advancing our understanding of the mechanisms underlying social behaviors across species.

## Methods

### Ethics statement

All experimental procedures were conducted in accordance with guidelines approved by our institutional Animal Care Committee and adhered to all Canadian Council on Animal Care regulations.

### Subjects and Housing

Subjects were 35 wild-type zebrafish (*Danio rerio*) bred in-house, 180 Pachón cavefish (*Astyanax mexicanus*) acquired from a local supplier (Tropical Fish Room, Brantford, ON), and 120 surface-dwelling *Astyanax mexicanus*, also bred in-house. In Experiment 1, we tested 35 zebrafish, 35 blind AM, and 30 surface AM to assess schooling behavior. Experiment 2 involved 55 blind AM reused from Experiment 4 to investigate the effects of hunger on schooling. Experiment 3 tested 90 blind AM and 90 surface fish to evaluate the impact of hormone administration on schooling behavior. Experiment 4 involved 55 blind AM and focused on the effects of increased dosages of certain hormones. With the exception of Experiment 2, each fish was only tested once. Fish in Experiment 2 had completed experiment 4 at least one month earlier.

Zebrafish were housed in 10-liter tanks in a high-density rack system (Pentair), with no more than 10 fish per tank, while blind and surface-dwelling AM were housed in groups of 5 to 20 in 10-gallon tanks (50 x 25 x 30 cm). All tanks were maintained at 23 ± 1°C with a 12 h:12 h light-dark cycle (lights on at 7:00 am). Water quality parameters, including pH, salinity, nitrate, and nitrite levels, were monitored daily. All fish were fed dried brine shrimp daily *ad lib*, except during Experiment 2, where fish were subjected to varied feeding schedules to manipulate nutritional state.

### Experimental Setup

Each experiment was conducted in a featureless, white circular tank (60 cm diameter; Figure S1), filled with 10 cm of water at 23 ± 1°C. The water in the experimental tank was changed daily, with temperature and salinity matched to those of the housing tanks. Behavioral trials were recorded using a video camera (Canon Vixia HF R700) mounted directly above the tank.

### Procedure

In all experiments, groups of fish (N = 5) were first gently netted from their home tanks into a bucket filled with home-tank water. Fish were then either transferred directly to the testing arena (Experiments 1 and 2) or given injections before testing (Experiments 3 and 4; see below). Fish were gently netted and released into the center of the experimental tank and were recorded swimming freely for 10 minutes (Videos S1 and S2). At the end of each trial, fish were gently netted back into the bucket and returned to their home tanks.

In Experiment 1, seven groups of zebrafish, seven groups of blind AM, and six groups of surface fish were tested to assess baseline shoaling behavior in each group. In Experiment 2, eleven groups of blind AM were randomly divided into two conditions. Six groups were food-deprived for 24 hours before testing (fasted condition), and five groups were fed *ad lib* 10 minutes before the testing (fed condition). To minimize the number of fish used, data from the unmanipulated blind AM in Experiment 1, which were fed *ad lib* approximately 3 to 5 hours before testing, were used as the control (post-absorption condition).

In Experiment 3, we tested the effects of social hormones on shoaling behavior in both blind and sighted AM. Subjects were randomly selected from their home tanks and assigned to one of five treatment groups immediately prior to testing, where they received an injection of either isotocin (IT; Carbetocin acetate, *MilliporeSigma Canada*), AVT ([Arg⁸]-Vasotocin, *VWR International*), an IT antagonist (L-368,899 hydrochloride, *Cayman Chemical*), an AVT antagonist (Manning Compound, *VWR International*), or saline (0.9% physiological saline).

Before injection, each fish was weighed by being placed into a beaker of water of known weight, and injection volumes were then calculated individually. Fish were positioned upside-down in a slit within a damp sponge for injection. Drugs were administered via intraperitoneal injection using a 10 µL, 26-gauge syringe (Hamilton 701N). The entire procedure took approximately 30 seconds per fish. Each group of five fish was injected sequentially and then transferred to a recovery tank for 10 minutes before testing.

All drugs were administered at 10 μg/g body weight, with injection volumes between 2 and 8 μL. For blind AM, five groups received saline, three received IT (denoted IT+), four received AVT (AVT+), three received the IT antagonist (IT−), and three received the AVT antagonist (AVT−). For surface fish, three groups received saline, four received IT+, three received AVT+, four received IT−, and four received AVT−.

In Experiment 4, we tested the effects of higher hormone dosages in blind AM only. Fish were randomly selected from their home tanks and assigned to one of four treatment groups immediately prior to testing, receiving either 20 μg/g or 40 μg/g doses of AVT+ or IT-following the same procedures as Experiment 3. Three groups received 20 μg/g of AVT, three received 40 μg/g of AVT, and five received 40 μg/g of IT−. Data from the saline-injected groups in Experiment 3 were used as a control. Our dosages were informed by previous research on social hormone administration in small fish species (116–118,142), as no previous studies have manipulated IT and AVT in AM.

### Analysis

Fish movements were tracked from video recordings using an automated tracking system (*IDTracker*; 143), which provided the precise location of each fish in each frame of the video. Using a custom script in R (144), data were processed to extract several standard metrics of collective movement for each frame: the inter-individual distance (IID), defined as the mean distance between an individual and all others, averaged across all fish; nearest-neighbor distance (NND), defined as the distance between each fish and its nearest neighbor, averaged over all individuals; and polarization, measuring the degree to which individuals are oriented in the same direction. These metrics have been validated in studies examining fish shoaling behavior and are widely recognized as robust indicators of group cohesion and alignment (5,145). All distance measurements were converted from pixels to centimeters using a scaling factor derived from the arena’s known diameter of 60 cm in the video.

To assess whether fish were actively shoaling or ignoring each other, we created a null model in R to simulate random movement. This model created 10,000 iterations of random point distributions (five points per iteration) within the experimental arena, calculating IID, NND, and polarization values for each configuration. These randomly generated values served as a baseline for comparison with the observed experimental data.

All statistical analyses were performed in *Mathematica* (v.12.0, *Wolfram Technologies*). Analysis of variance (ANOVA) tests were used to evaluate significant differences between species, experimental conditions, and hormonal treatments. Specifically, one-way ANOVAs were applied to assess differences in IID, NND, and polarization between treatment groups or species. Two-way ANOVAs were used to examine interactions between species (blind AM, surface-dwelling AM, and zebrafish) and treatment conditions (hormonal treatments, nutritional state). When the ANOVA indicated significant effects, post-hoc tests were conducted, and a Bonferroni correction was applied to all tests. Comparisons of a group to the theoretical null model were conducted using T-tests. We additionally report the power of each test, using η^2^ for ANOVAs and Cohen’s D for T-tests.

Given that we tracked our fish in every frame of the video (30 times per second), the values of all our measures are not independent across measures, which violates the assumptions of our analysis methods. To address this, before conducting any analyses, we down-sampled our data, taking just one frame from each minute of each trial (i.e., we took 10 evenly spaced frames from the data for each trial), ensuring statistical validity by reducing autocorrelation in the dataset.

All the data reported here are available on our OSF repository, at https://osf.io/9jky4/?view_only=c8e10cd8b43c4e58b3d84d0de6f0d876 [This repository will be made public upon publication].

## Results

### Experiment 1

We found significant differences between all species or strains on all measures of shoaling behavior (Figure 1 A-C. Means ± SD in Table S1, post-hoc tests in Table S2. IID: F(2,197) = 1216.4, p < 0.00001, η^2^ = 0.93; NND: F(2,197) = 927.7, p < 0.00001, η^2^ = 0.90; Polarization: F(2,197) = 233.1, p < 0.00001, η^2^ = 0.70). We additionally compared the experimental data to our null distribution to test the assumption that blind AM were simply ignoring each other in the tank (means ± SD in Table S1). We found that both zebrafish and sighted AM shoaled more tightly than predicted by the model (NND: zebrafish, T(52.7) = 5.13, p < 0.00001, Cohen’s D = 1.11; sighted AM, T(50.9) = 6.90, p < 0.00001, D = 1.44. IID: zebrafish, T(62.1) = 6.97, p < 0.00001, D = 1.45; sighted AM, T(54.4) = 10.75, p < 0.00001, D = 2.23) but blind AM maintained *greater* distances between individuals than predicted by the model (NND: T(53.8) = −3.8, p = 0.0004, D = 0.82. IID: T(58.1) = −8.9, p < 0.00001, D = 1.88). Zebrafish and sighted AM groups were more polarized than the model (zebrafish: T(67.3) = −8.44, p < 0.00001, D = 1.72; sighted AM: T(71.5) = −10.0, p < 0.00001, D = 2.02) but there was no difference in polarization between the model and blind AM (T(56.8) = 0.26, p = 0.80, D = 0.05).

**Figure 1.**
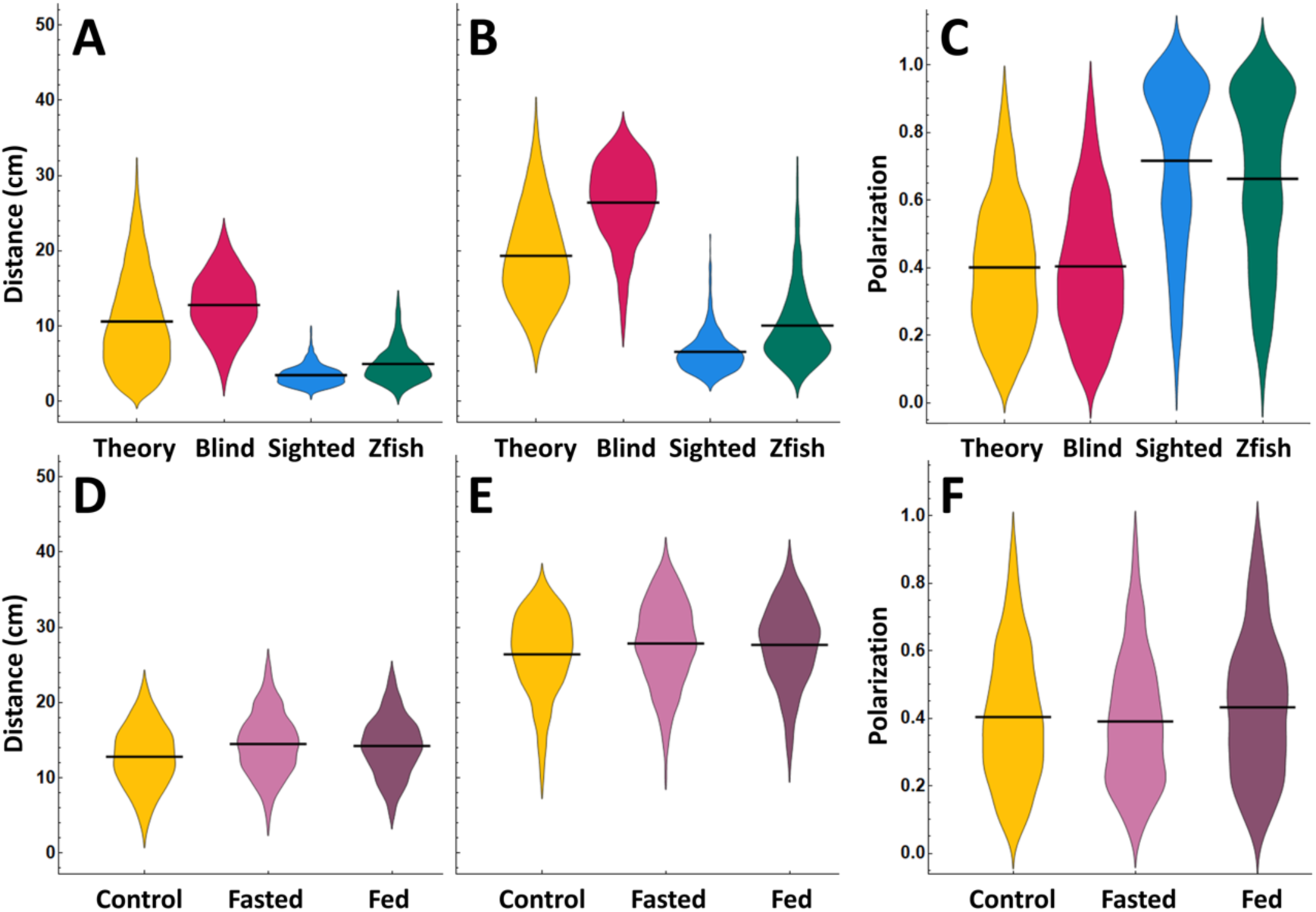
Results of Experiments 1 and 2. Distribution plots of NND (A, D), IID (B, E), and polarization (C, F) for the different species/strains in Experiment 1 (A-C) and the different feeding conditions in Experiment 2 (D-F). Horizontal black lines indicate the mean of each distribution.

Distributions of polarization in both zebrafish and sighted AM (Figure 1C) displayed a characteristic bimodal profile, indicating the presence of two kinds of shoaling (sometimes referred to as schooling and shoaling; 146), but no such distinction was evident in the polarizations of blind AM.

### Experiment 2

Hunger levels had a significant impact on shoaling behavior in blind cavefish (Figure 1 D-F). Both hungry and recently fed fish schooled less tightly than control fish (means ± SD in Table S1; post-hocs in Table S3. IID: F(2,177) = 10.34, p = 0.00006, η^2^ = 0.10; NND: F(2,177) = 29.57, p < 0.00001, η^2^ = 0.25). Polarization also differed significantly across conditions (F(2,177) = 7.05, p = 0.001, η^2^ = 0.07), with fed fish showing higher polarization than hungry fish (Table S3).

### Experiment 3

We found a significant main effect of strain and an interaction between strain and drug treatment but no main effect of drug treatment on both IID (Figure 2. Means ± SD in Table S1; post-hocs in Tables S4-S5. Strain: F(1, 350) = 6712, p < 0.00001, η^2^ = 0.94; drug: F(4, 350) = 2.78, p = 0.03, η^2^ = 0.002; strain x drug: F(4, 350) = 16.79, p < 0.00001, η^2^ = 0.01) and NND (strain: F(1,350) = 6547, p < 0.00001, η^2^ = 0.94; drug: F(4,350) = 1.60, p = 0.17, η^2^ = 0.001; strain x drug: F(4,350) = 11.71, p < 0.00001, η^2^ = 0.007). On polarization, we found main effects of both strain and drug, as well as an interaction (strain: F(1,350) = 650, p < 0.00001, η^2^ = 0.59; drug: F(4,350) = 15.0, p < 0.00001, η^2^ = 0.05; strain x drug: F(4,350) = 10.04, p < 0.00001, η^2^ = 0.04).

**Figure 2.**
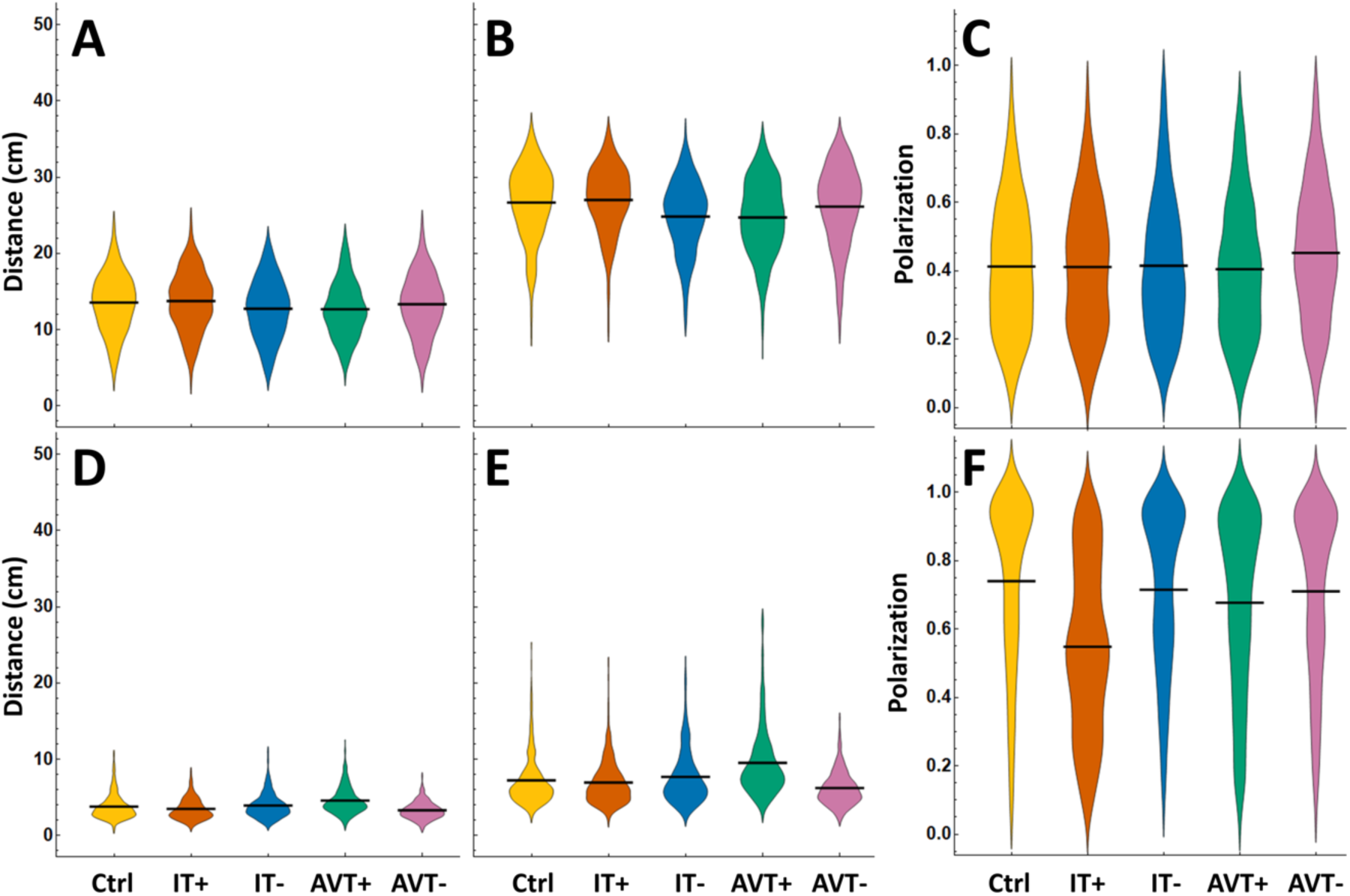
Results of experiment 3. Distribution plots of NND (A, D), IID (B, E), and polarization (C, F) for the blind (A-C) and sighted AM (D-F). Ctrl = control, IT+ = isotocin, IT−= isotocin antagonist, AVT+ = vasotocin, AVT−= vasotocin antagonist. Horizontal black lines indicate the mean of each distribution.

### Experiment 4

The administration of higher AVT+ doses (at 20 and 40 μg/g) in blind AM resulted in significantly looser shoals compared to the control and 10 μg/g doses (Figure 3. Means ± SD in Table S1; post-hocs in Table S6. IID: F(3,146) = 15.49, p < 0.00001, η^2^ = 0.24; NND: F(3,146) = 15.19, p < 0.00001, η^2^ = 0.24), that were also slightly less polarized (F(3,146) = 4.28, p = 0.006, η^2^ = 0.08). Increased doses of IT−(40 μg/g) led to significantly lower IID compared to the control group (F(2,127) = 13.36, p < 0.00001, η^2^ = 0.17), but not NND (F(2,127) = 3.94, p = 0.022, η^2^ = 0.06), and had no effect on polarization (F(2,127) = 1.49, p = 0.23, η^2^ = 0.02).

**Figure 3.**
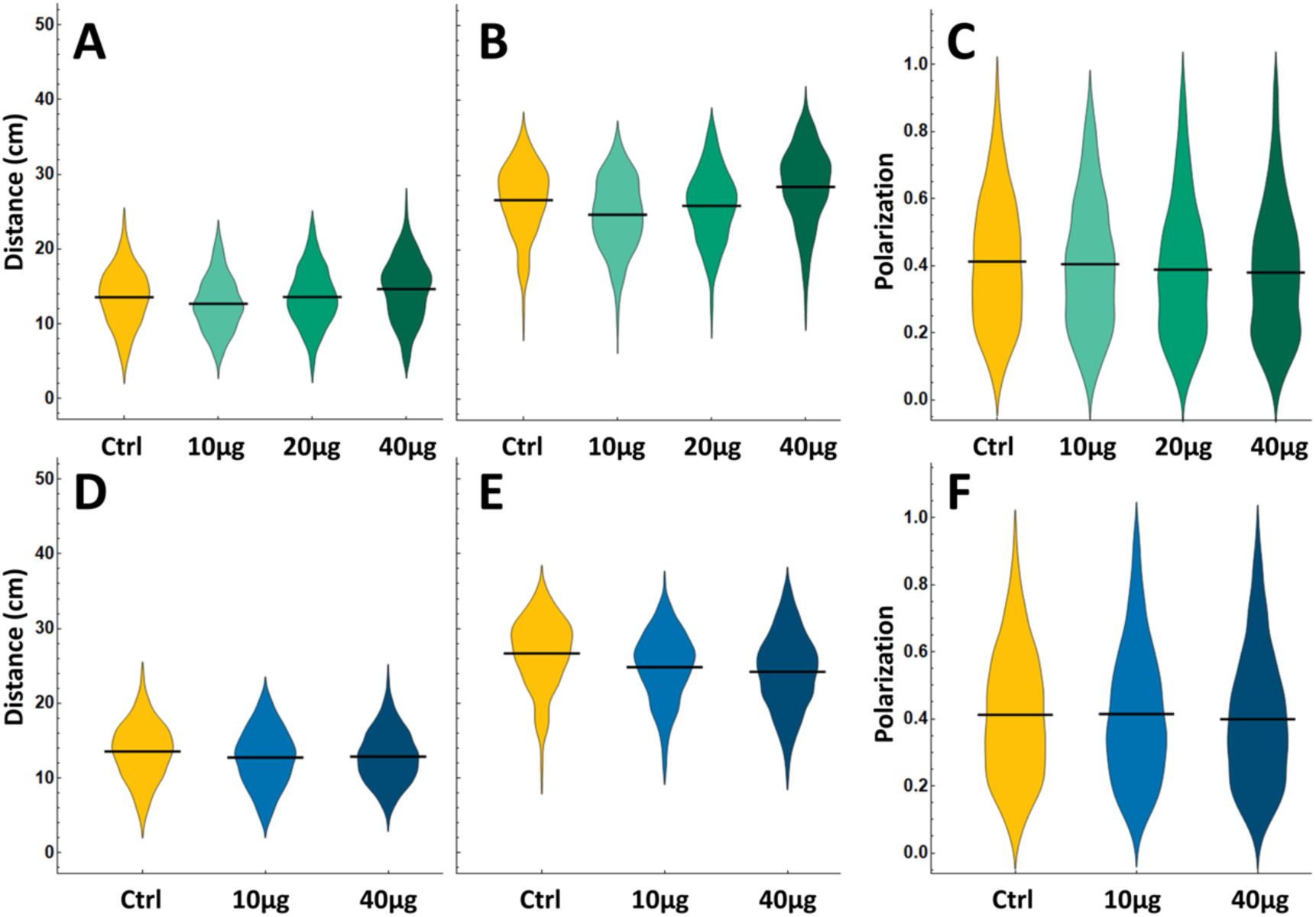
Results of experiment 4. Distribution plots of NND (A, D), IID (B, E), and polarization (C, F) for increasing concentrations of vasotocin (A-C) and an isotocin antagonist (D-F). Horizontal black lines indicate the mean of each distribution.

## Discussion

Our study compared the shoaling behaviors of blind *Astyanax mexicanus* (AM) with their surface-dwelling conspecifics and zebrafish, examining how nutritional state and neurohormonal modulation influence these behaviors. These findings provide new insights into how environmental pressures, sensory adaptations, and neuroendocrine regulation shape social behavior in species adapted to extreme environments.

We found, unsurprisingly, that both sighted AM and zebrafish formed cohesive shoals, maintaining close distances—much closer than would be expected by chance (Figure 1 A-B; Video S2)—and displayed coordinated movement, as evidenced by high polarization values and bimodal distributions characteristic of schooling behavior (Figure 1C; 146). Blind AM, in contrast, neither coordinated their movements—as the polarization of the group was identical to what would be expected by chance—nor formed shoals. Notably, they maintained significantly *greater* distances between individuals than would be expected if they were simply ignoring each other, suggesting active avoidance rather than social indifference (Video S1).

We also found that both food deprivation and feeding increased the distances that blind AM maintained from one another (Figure 1 D-E). However, groups of recently fed fish were more polarized than fasted groups (Figure 1F). Blind AM also moved closer together when we increased AVT levels or blocked the action of IT in their brains (Figure 2 A-B), though the former effect was reversed at higher concentrations of AVT (Figure 3 A-B). Interestingly, AVT had the opposite effect on sighted AM, moving them further apart (Figure 2 D-E). Increasing IT levels caused a drastic reduction in the polarization of sighted AM groups, and the polarization of blind AM groups varied inversely with AVT: decreasing AVT increased polarization (Figure 2C), while high doses of AVT reduced it (Figure 3C).

Our findings indicate that altering nutritional state or manipulating levels of IT and AVT in blind AM can alter their group cohesion and polarization. Although the specific directions and magnitudes of these effects were not always as predicted (see below), the mere fact that hunger and hormonal changes impact shoal density suggests that blind AM’s lack of shoaling behavior stems more from a reduced motivation to aggregate than from an inability to detect relative locations of conspecifics. This conclusion is further strengthened by our finding that groups of unmanipulated blind AM do not simply ignore one another but rather actively avoid each other, directing their movements based on the positions of others but being repulsed from rather than attracted to the, (as shoaling fish would be). Together, these results provide valuable insights into the evolutionary pressures that shape social behaviors.

While blind AM retain the ability to perceive their surroundings through their enhanced lateral line system, our results suggest that their motivation to engage in social behaviors like shoaling has been lost since they split from the sighted morph approximately 20,000 years ago. This behavioral shift likely reflects adaptation to cave environments where aggregation is disadvantageous due to resource scarcity and the absence of predators, as close grouping could increase risks of kleptoparasitism and competition, making solitary swimming more advantageous (101). Similar shifts away from social behaviors have been observed in other species experiencing changes in predation pressures, including guppies (29). Solitary swimming may also allow the lateral line system to detect subtle water movements with greater precision by reducing interference from conspecifics, thus enhancing navigation and foraging in darkness (60, 81,85,148). While blind AM have often been described as ‘asocial’ (147), the term ‘anti-social’ may better capture their active avoidance of conspecifics.

Our results also reveal that nutritional state modulates social behavior in blind AM, which are constantly food-oriented. Contrary to our initial expectations, both the fasted and fed groups maintained greater distances from each other than the post-absorptive control group (Figure 1 D-E). This pattern suggests that both hunger and recent feeding may shift behavioral priorities toward foraging or digestion. Such behavior likely reflects an energy-conserving adaptation, advantageous in environments where food resources are both scarce and unpredictable.

The social behaviors we observed in surface-dwelling AM under increased AVT are consistent with findings in other social fish species, where AVT administration tended to promote social distancing or even aggression rather than cohesion (109,113,114,116,125). These findings align with AVT’s known role in enhancing territoriality and defensive behavior across taxa.

A particularly striking result is our finding that increasing the amount of AVT in their brains increased cohesion in groups of blind AM, but decreased it in sighted AM. This, together with the reduced density of AVT-producing cells in the blind AM brain (94), provides strong evidence that this molecule and its receptors have been the targets of strong selection since the two strains diverged. However, we also found that higher AVT doses reversed the effect in blind AM, leading to reduced cohesion. The higher AVT doses may have been necessary to elicit a comparable response to that seen in sighted AM with lower doses, suggesting an altered sensitivity to AVT in the blind morph. In either case, as AVT is closely involved in social cognition in a wide range of species, changes in the AVT system may largely be responsible for having reversed how blind AM react to conspecifics, replacing attraction with repulsion (assuming the sighted form is ancestral).

The role of IT in modulating social behavior also differed notably between the two morphs. In blind AM, blocking IT receptors with an antagonist—at both low and high doses— decreased distances between individuals, partially restoring shoaling-like behavior. This suggests that IT typically functions to suppress social cohesion in blind AM (Figure 2 A-B). No such effect was observed in sighted fish; however, increasing IT levels decreased their polarization. These findings stand in contrast to research on other fish species, such as guppies and goldfish, as well as studies on OT in mammals, where this hormone often enhances social bonding, facilitating group cohesion and affiliation (132–139).

However, our results from blind AM align with those seen in male *N. pulcher*, where IT administration inhibits grouping behavior, while IT antagonists have been found to increase group cohesion (122). These cichlids have higher levels of IT and lower levels of AVT than closely related less social species (124,149). Although their social behaviors contrast (blind AM are solitary while *N. pulcher* are social), their neurohormonal responses to IT manipulation, as well as their baseline hormone levels, reveal notable similarities. This may suggest that, despite divergent ecological demands, certain neuroendocrine systems may share some evolutionary pathways, shaping distinct social strategies through similar mechanisms.

One possible explanation for the IT antagonist’s increasing proximity in blind AM is the social salience hypothesis, which suggests that OT/IT may increase the salience of social stimuli, both positive and negative (128–130). In the case of blind AM, where shoaling may be maladaptive, social stimuli may be perceived as aversive, leading them to actively avoid conspecifics. Blocking IT, however, may reduce their sensitivity to these aversive social signals, leading to an increase in proximity, as we observed in our IT-antagonized blind AM groups.

## Conclusion

Neuropeptides such as AVP/AVT and OT/IT play a complex role in shaping social behaviors across taxa. Our findings reveal that the solitary nature of blind AM is not due to an inability to detect conspecifics but rather reflects a decreased motivation to maintain social proximity. This highlights how blind AM have adapted their social behavior to the unique conditions of cave environments where predators are absent and competition for scarce resources favors solitary foraging over group living (53). The contrasting responses to AVT in blind and surface AM suggest that changes in the AVT system may have played a major role in restructuring social behavior since these morphs diverged. AVT enhances social cohesion in blind AM, whereas it promotes social distancing in the sighted morph, as it does in other social fish species (109,125).

These findings not only shed light on the evolution of social behavior in cavefish but also have broader implications for understanding how environmental pressures shape sociality through neurohormonal pathways. Our ability to modulate social behaviors through hormone administration suggests that similar mechanisms might influence the evolution of new social dynamics in other species experiencing drastic ecological changes, including those driven by human activity. For instance, intense fishing pressures can select against shoaling behaviors in marine populations, leading to social shifts that mirror those seen in blind AM (150). Further exploration of the adaptability of neurohormonal systems will improve our ability to predict behavioral responses to shifting ecosystems, including those impacted by human activity, ultimately aiding conservation efforts and the management of species under pressure.

## References

1. Wisenden BD. Factors affecting mate desertion by males in free-ranging convict cichlids (*Cichlasoma nigrofasciatum*). Behav Ecol. 1994;5(4):439–47. 10.1093/beheco/5.4.439

2. Wong M, Balshine S. The evolution of cooperative breeding in the African cichlid fish, *Neolamprologus pulcher*. Biol Rev Camb Philos Soc. 2011;86(4):511–30. 10.1111/j.1469-185X.2010.00158.x

3. Fox HE, White SA, Kao MH, Fernald RD. Stress and dominance in a social fish. J Neurosci. 1997;17(16):6463–9. 10.1523/JNEUROSCI.17-16-06463.1997

4. Dugatkin LA, Mesterton-Gibbons M. Cooperation among unrelated individuals: reciprocal altruism, by-product mutualism and group selection in fishes. BioSystems. 1996;37(2-3):19–30. 10.1016/0303-2647(95)01542-6

5. Miller N, Gerlai R. Quantification of shoaling behaviour in zebrafish (*Danio rerio*). Behav Brain Res. 2007;184(2):157–66. 10.1016/j.bbr.2007.07.007

6. Gautrais J, Ginelli F, Fournier R, Blanco S, Soria M, Chaté H, Theraulaz G. Deciphering interactions in moving animal groups. PLoS Comput Biol. 2012;8(9). 10.1371/journal.pcbi.1002678

7. Lopez U, Gautrais J, Couzin ID, Theraulaz G. From behavioural analyses to models of collective motion in fish schools. Interface Focus. 2012;2(6):693–707. 10.1098/rsfs.2012.0033

8. Gerlai R. Social behavior of zebrafish: From synthetic images to biological mechanisms of shoaling. J Neurosci Methods. 2014;234:59–65. 10.1016/j.jneumeth.2014.04.028

9. Magurran AE, Oulton WJ, Pitcher TJ. Vigilant behavior and shoal size in minnows. Z Tierpsychol. 1985;67(1-4):167–78. 10.1111/j.1439-0310.1985.tb01386.x

10. Godin JGJ, Classon LJ, Abrahams MV. Group vigilance and shoal size in a small characin fish. Behaviour. 1988;104(1-2):29–40. 10.1163/156853988X00584

11. Ward AJW, Herbert-Read JE, Sumpter DJT, Krause J. Fast and accurate decisions through collective vigilance in fish shoals. Proc Natl Acad Sci USA. 2011;108(6):2312–5. 10.1073/pnas.1007102108

12. Ward AJW, Webster MM. Sociality: The behaviour of group living animals. Springer; 2016. 10.1007/978-3-319-28585-6

13. Krause J, Ruxton GD. Living in groups. Oxford University Press; 2002.

14. Ruxton GD, Jackson AL, Tosh CR. Confusion of predators does not rely on specialist coordinated behavior. Behav Ecol. 2007;18(3):590–6. 10.1093/beheco/arm009

15. Landeau L, Terborgh J. Oddity and the confusion effect in predation. Anim Behav. 1986;34:1372–80. 10.1016/s0003-3472(86)80208-1

16. Larsson M. Possible functions of the octavolateralis system in fish schooling. Fish Fish. 2009;10(3):344–53. 10.1111/j.1467-2979.2009.00330.x

17. Roberts G. Why individual vigilance declines as group size increases. Anim Behav. 1996;51:1077–86. 10.1006/anbe.1996.0109

18. Day RL, MacDonald T, Brown C, Laland KN, Reader SM. Interactions between shoal size and conformity in guppy social foraging. Anim Behav. 2001;62(5):917–25. 10.1006/anbe.2001.1820

19. Baird TA, Ryer CH, Olla BL. Social enhancement of foraging on an ephemeral food source in juvenile walleye pollock (Theragra chalcogramma). Environ Biol Fishes. 1991;31(3):307–11. 10.1007/bf00000697

20. Magurran AE. The adaptive significance of schooling as an anti-predator defence in fish. Ann Zool Fenn. 1990;27:51–66.

21. Partridge BL. The structure and function of fish schools. Sci Am. 1982;246(6):114–23. 10.1038/scientificamerican0682-114

22. Pitcher TJ, Magurran AE, Winfield IJ. Fish in larger shoals find food faster. Behav Ecol Sociobiol. 1982;10(2):149–51. 10.1007/BF00300175

23. Smith MFL, Warburton K. Predator shoaling moderates the confusion effect in blue-green chromis, *Chromis viridis*. Behav Ecol Sociobiol. 1992;30:103–7. 10.1007/BF00173946

24. Ranta E, Rita H, Lindström K. Competition versus cooperation: Success of individuals foraging alone and in groups. Am Nat. 1993;142(1):42–58. 10.1086/285528

25. Thünken T, Landmann J, Bakker TCM, Baldauf SA. Increased levels of perceived competition decrease juvenile kin-shoaling preferences in a cichlid fish. Am Nat. 2020;195(1):1–11. 10.1086/707747

26. Ioannou CC. Swarm intelligence in fish? The difficulty in demonstrating distributed and self-organised collective intelligence in (some) animal groups. Behav Process. 2017;141(Pt 2):141–51. 10.1016/j.beproc.2016.10.005

27. Kasumyan AO, Pavlov DS. On the problem of the evolutionary origin of schooling behavior of fish. J Ichthyol. 2023;63(6):1374–89. 10.1134/S0032945223070135

28. Blumstein DT, Daniel JC. The loss of anti-predator behavior following isolation on islands. Proc Biol Sci. 2005;272(1573):1663–8. 10.1098/rspb.2005.3147

29. Ioannou CC, Ramnarine IW, Torney CJ. High-predation habitats affect the social dynamics of collective exploration in a shoaling fish. Sci Adv. 2017;3(5). 10.1126/sciadv.1602682

30. Herbert-Read JE, Wade ASI, Ramnarine IW, Ioannou CC. Collective decision-making appears more egalitarian in populations where group fission costs are higher. Biol Lett. 2019;15(12):20190556. 10.1098/rsbl.2019.0556

31. Magurran AE, Seghers BH, Carvalho GR, Shaw PW. Behavioural consequences of an artificial introduction of guppies (*Poecilia reticulata*) in N. Trinidad: evidence for the evolution of anti-predator behaviour in the wild. Proc Biol Sci. 1992;248(1322):117–22. 10.1098/rspb.1992.0050

32. Seghers BH. Schooling behavior in the guppy (*Poecilia reticulata*): An evolutionary response to predation. Evolution. 1974;28(3):486–9. 10.2307/2407174

33. Huizinga M, Ghalambor CK, Reznick DN. The genetic and environmental basis of adaptive differences in shoaling behaviour among populations of Trinidadian guppies, *Poecilia reticulata*. J Evol Biol. 2009;22(9):1860–6. 10.1111/j.1420-9101.2009.01799.x

34. Song Z, Boenke MC, Rodd FH. Interpopulation differences in shoaling behaviour in guppies (*Poecilia reticulata*): Roles of social environment and population origin. Ethology. 2011;117(11):1009–18. 10.1111/j.1439-0310.2011.01952.x

35. Wade ASI, Ramnarine IW, Ioannou CC. The effect of group size on the speed of decision making depends on compromise and predation risk across populations in the guppy *Poecilia reticulata*. Behaviour. 2020;157(14-15):1173–92. 10.1163/1568539X-bja10044

36. Stockwell CA, Schmelzer MR, Gillis BE, Anderson CM, Wisenden BD. Ignorance is not bliss: Evolutionary naiveté in an endangered desert fish and implications for conservation. Proc Biol Sci. 2022;289(1978):20220752. 10.1098/rspb.2022.0752

37. Webber HM, Haines TA. Mercury effects on predator avoidance behavior of a forage fish, golden shiner (*Notemigonus crysoleucas*). Environ Toxicol Chem. 2009;22(7):1556–61. 10.1002/etc.5620220718

38. Scott GR, Sloman KA. The effects of environmental pollutants on complex fish behaviour: Integrating behavioural and physiological indicators of toxicity. Aquat Toxicol. 2004;68(4):369–92. 10.1016/j.aquatox.2004.03.016

39. Mao X. Review of fishway research in China. Ecol Eng. 2018;115:91–5. 10.1016/j.ecoleng.2018.01.010

40. Herbert-Read JE, Kremer L, Bruintjes R, Radford AN, Ioannou CC. Anthropogenic noise pollution from pile-driving disrupts the structure and dynamics of fish shoals. Proc Biol Sci. 2017;284(1868):20171627. 10.1098/rspb.2017.1627

41. Rodriguez-Pinto II, Rieucau G, Handegard NO, Boswell KM. Environmental context elicits behavioural modification of collective state in schooling fish. Anim Behav. 2020;167:75–84. 10.1016/j.anbehav.2020.05.002

42. Mukherjee I, Bhat A. The impact of predators and vegetation on shoaling in wild zebrafish. R Soc Open Sci. 2024;11(9):240760. 10.1098/rsos.240760

43. Krause J, Hartmann N, Pritchard VL. The influence of nutritional state on shoal choice in zebrafish, *Danio rerio*. Anim Behav. 1999;57(4):771–5. 10.1006/anbe.1998.1010

44. Hensor EMA, Magurran AE, Pitcher TJ. Shoaling and foraging in the banded killifish (*Fundulus diaphanus*): The effects of nutritional state and shoal size. Anim Behav. 2003;66(3):463–70. 10.1006/anbe.2003.2239

45. Sogard SM, Olla BL. The influence of hunger and predation risk on group cohesion in a pelagic fish, walleye pollock (*Theragra chalcogramma*). Environ Biol Fishes. 1997;50(4):405–13. 10.1023/A:1007376115374

46. Krause J. The influence of hunger on shoal size choice by three-spined sticklebacks, *Gasterosteus aculeatus*. J Fish Biol. 1993;43(5):775–80. 10.1111/j.1095-8649.1993.tb01154.x

47. Aimon C, Le Bayon N, Le Floch S, Claireaux G. Food deprivation reduces social interest in the European sea bass (*Dicentrarchus labrax*). J Exp Biol. 2019;222(3). 10.1242/jeb.190553

48. Fumey J, Hinaux H, Noirot C, Thermes C, Rétaux S, Casane D. Evidence for late pleistocene origin of *Astyanax mexicanus* cavefish. BMC Evol Biol. 2018;18(1):1156–71. 10.1186/s12862-018-1156-7

49. Herman A, Brandvain Y, Weagley J, et al. The role of gene flow in rapid and repeated evolution of cave-related traits in Mexican tetra, *Astyanax mexicanus*. Mol Ecol. 2018;27(22):4397–416. 10.1111/mec.14877

50. Parzefall J. A review of morphological and behavioural changes in the cave molly, *Poecilia mexicana*, from Tabasco, Mexico. Environ Biol Fishes. 2001;62(1-3):263–75. 10.1023/A:1011899817764

51. Mitchell RW, Russell WH, Elliott WR. Mexican eyeless characin fishes, genus *Astyanax*: Environment, distribution, and evolution. KIP Monographs. 1977;(17). University of South Florida. Available from: https://digitalcommons.usf.edu/kip_monographs/17

52. Espinasa L, Ornelas-García CP, Legendre L, et al. Discovery of two new *Astyanax* cavefish localities leads to further understanding of the species biogeography. Diversity. 2020;12(10). 10.3390/d12100368

53. Parzefall J. Field observation in epigean and cave populations of Mexican characid *Astyanax mexicanus* (Pisces, Characidae). Mem Biospeol. 1983;10:171–6.

54. Jeffery WR. Cavefish as a model system in evolutionary developmental biology. Dev Biol. 2001;231(1):1–12. 10.1006/dbio.2000.0121

55. Jeffery WR. Regressive evolution in *Astyanax* cavefish. Annu Rev Genet. 2009;43:25–47. 10.1146/annurev-genet-102108-134216

56. Elmer KR, Meyer A. Adaptation in the age of ecological genomics: Insights from parallelism and convergence. Trends Ecol Evol. 2011;26(6):298–306. 10.1016/j.tree.2011.02.008

57. Aspiras ACR, Rohner N, Martineau B, Borowsky RL, Tabin CJ. Melanocortin 4 receptor mutations contribute to the adaptation of cavefish to nutrient-poor conditions. Proc Natl Acad Sci USA. 2015;112(31):9668–73. 10.1073/pnas.1510802112

58. Agnès F, Torres-Paz J, Michel P, Rétaux S. A 3D molecular map of the cavefish neural plate illuminates eye-field organization and its borders in vertebrates. Development. 2022;149. 10.1242/dev.199966

59. Duboué ER, Keene AC, Borowsky RL. Evolutionary convergence on sleep loss in cavefish populations. Curr Biol. 2011;21(8):671–6. 10.1016/j.cub.2011.03.020

60. Elipot Y, Hinaux H, Callebert J, Rétaux S. Evolutionary shift from fighting to foraging in blind cavefish through changes in the serotonin network. Curr Biol. 2013;23(1):1–10. 10.1016/j.cub.2012.10.044

61. Elipot Y, Hinaux H, Callebert J, Launay JM, Blin M, Rétaux S. A mutation in the enzyme monoamine oxidase explains part of the *Astyanax* cavefish behavioral syndrome. Nat Commun. 2014;5:3647. 10.1038/ncomms4647

62. Hyacinthe C, Attia J, Rétaux S. Evolution of acoustic communication in blind cavefish. Nat Commun. 2019;10:4231. 10.1038/s41467-019-12201-9

63. Hinaux H, Devos L, Blin M, et al. Sensory evolution in blind cavefish is driven by early embryonic events during gastrulation and neurulation. Development. 2016;143(24):4521–32. 10.1242/dev.141291

64. Kowalko JE, Rohner N, Linden TA, Rompani SB, Warren WC. Loss of schooling behavior in cavefish through sight-dependent and sight-independent mechanisms. Curr Biol. 2013;23(19):1874–83. 10.1016/j.cub.2013.07.056

65. Torres-Paz J, Hyacinthe C, Pierre C, Rétaux S. Towards an integrated approach to understand Mexican cavefish evolution. Biol Lett. 2018;14(8):20180101. 10.1098/rsbl.2018.0101

66. Varatharasan N, Croll RP, Franz-Odendaal T. Taste bud development and patterning in sighted and blind morphs of *Astyanax mexicanus*. Dev Dyn. 2009;238(12):3056–64. 10.1002/dvdy.22128

67. Parzefall J. On the heredity of behavior patterns in cave animals and their epigean relatives. Bull Nat Speleol Soc. 1985;47:128–35.

68. Avise JC, Selander RK. Evolutionary genetics of cave-dwelling fishes of the genus *Astyanax*. Evolution. 1972;26(1):1–19. 10.1111/j.1558-5646.1972.tb00170.x

69. Wilkens H. Evolution and genetics of epigean and cave *Astyanax fasciatus* (Characidae, Pisces). In: Evolutionary Biology. Springer; 1988. p. 271–367. 10.1007/978-1-4613-1043-3_8

70. Hüppop K. Food-finding ability in cave fish (*Astyanax fasciatus*). Int J Speleol. 1987;16(1):59–66. 10.5038/1827-806X.16.1.4

71. Rodríguez-Morales R. Sensing in the dark: Constructive evolution of the lateral line system in blind populations of *Astyanax mexicanus*. Ecol Evol. 2024;14(4). 10.1002/ece3.11286

72. Wilkens H, Strecker U. Evolution in the dark: Darwin’s loss without selection. Springer Nature; 2017.

73. Krishnan J, Rohner N. Cavefish and the basis for eye loss. Philos Trans R Soc Lond B Biol Sci. 2017;372(1713):20150487. 10.1098/rstb.2015.0487

74. Dijkgraaf S. The functioning and significance of the lateral-line organs. Biol Rev Camb Philos Soc. 1963;38(1):51–105. 10.1111/j.1469-185X.1963.tb00654.x

75. Kalmijn AJ. Functional evolution of lateral line and inner ear sensory systems. In: Coombs S, Görner P, Münz H, editors. The mechanosensory lateral line. Springer; 1989. p. 187–215. 10.1007/978-1-4612-3560-6_9

76. Denton EJ, Gray JA. Some observations on the forces acting on neuromasts in fish lateral line canals. In: Coombs S, Görner P, Münz H, editors. The mechanosensory lateral line: Neurobiology and evolution. Springer; 1989. p. 229–46. 10.1007/978-1-4612-3560-6_11

77. Jeffery WR, Strickler AG, Guiney S, Heyser DG, Tomarev SI. Prox1 in eye degeneration and sensory organ compensation during development and evolution of the cavefish *Astyanax*. Dev Genes Evol. 2000;210(5):223–30. 10.1007/s004270050308

78. Teyke T. Collision with and avoidance of obstacles by blind cave fish *Anoptichthys jordani* (Characidae). J Comp Physiol A. 1985;157(6):837–43. 10.1007/BF01350081

79. Teyke T. Flow field, swimming velocity and boundary layer: parameters which affect the stimulus for the lateral line organ in blind fish. J Comp Physiol A. 1988;163(1):53–61. 10.1007/bf00611996

80. Teyke T. Morphological differences in neuromasts of the blind cave fish *Astyanax hubbsi* and the sighted river fish *Astyanax mexicanus*. Brain Behav Evol. 1990;35(1):23–30. 10.1159/000115853

81. Yoshizawa M, Gorički Š, Soares D, Jeffery WR. Evolution of a behavioral shift mediated by superficial neuromasts helps cavefish find food in darkness. Curr Biol. 2010;20(18):1631–6. 10.1016/j.cub.2010.07.017

82. Gertychowa R. Studies on the ethology and space orientation of the blind cave fish *Anoptichthys jordani* Hubbs et Innes 1936 (Characidae). Folia Biol (Krakow). 1970;18(1):9–69. PMID: 5433416.

83. John KR. Observations on the behavior of blind and blinded fishes. Copeia. 1957;1957(2):123–32. 10.2307/1439399

84. Windsor SP, Tan D, Montgomery JC. Swimming kinematics and hydrodynamic imaging in the blind Mexican cave fish (*Astyanax fasciatus*). J Exp Biol. 2008;211(18):2950–9. 10.1242/jeb.020453

85. Sharma S, Coombs S, Patton P, de Perera TB. The function of wall-following behaviors in the Mexican blind cavefish and a sighted relative, the Mexican tetra (*Astyanax*). J Comp Physiol A. 2009;195:225–40. 10.1007/s00359-008-0400-9

86. Patton P, Windsor S, Coombs S. Active wall following by Mexican blind cavefish (*Astyanax mexicanus*). J Comp Physiol A. 2010;196:853–67. 10.1007/s00359-010-0567-8

87. Hassan ES. Hydrodynamic imaging of the surroundings by the lateral line of the blind cave fish *Anoptichthys jordani*. In: Coombs S, Görner P, Münz H, editors. The Mechanosensory Lateral Line. Springer; 1989. p. 217–27. 10.1007/978-1-4612-3560-6_10

88. Yamamoto Y, Stock DW, Jeffery WR. Hedgehog signalling controls eye degeneration in blind cavefish. Nature. 2004;431(7010):844–7. 10.1038/nature02864

89. Yamamoto Y, Jeffery WR. Central role for the lens in cave fish eye degeneration. Science. 2000;289(5479):631–3. 10.1126/science.289.5479.631

90. Yoshizawa M, Yamamoto Y, O’Quin KE, Jeffery WR. Evolution of an adaptive behavior and its sensory receptors promotes eye regression in blind cavefish. BMC Biol. 2012;10(1):108. 10.1186/1741-7007-10-108

91. Yamamoto Y, Byerly MS, Jackman WR, Jeffery WR. Pleiotropic functions of embryonic sonic hedgehog expression link jaw and taste bud amplification with eye loss during cavefish evolution. Dev Biol. 2009;330(1):200–11. 10.1016/j.ydbio.2009.03.003

92. Menuet A, Alunni A, Joly JS, Jeffery WR, Rétaux S. Expanded expression of Sonic Hedgehog in *Astyanax* cavefish: Multiple consequences on forebrain development and evolution. Development. 2007;134(5):845–55. 10.1242/dev.02780

93. Pottin K, Hinaux H, Rétaux S. Restoring eye size in *Astyanax mexicanus* blind cavefish embryos through modulation of the Shh and Fgf8 forebrain organizing centres. Development. 2011;138(12):2467–76. 10.1242/dev.054106

94. Alié A, Devos L, Torres-Paz J, et al. Developmental evolution of the forebrain in cavefish, from natural variations in neuropeptides to behavior. eLife. 2018;7. 10.7554/eLife.32808

95. Yoshizawa M. Behaviors of cavefish offer insight into developmental evolution. Mol Reprod Dev. 2015;82(4):268–80. 10.1002/mrd.22471

96. Burchards H, Dölle A, Parzefall J. Aggressive behaviour of an epigean population of *Astyanax mexicanus* (Characidae, Pisces) and some observations of three subterranean populations. Behav Process. 1985;11(3):225–35. 10.1016/0376-6357(85)90017-8

97. Langecker TG, Neumann B, Hausberg C, Parzefall J. Evolution of the optical releasers for aggressive behavior in cave-dwelling *Astyanax fasciatus* (Teleostei, Characidae). Behav Process. 1995;34(2):161–7. 10.1016/0376-6357(94)00063-M

98. Espinasa L, Collins E, Ornelas García CP, et al. Divergent evolutionary pathways for aggression and territoriality in *Astyanax* cavefish. Subterr Biol. 2022;43:169–83. 10.3897/subtbiol.73.79318

99. Patch A, Paz A, Holt KJ, et al. Kinematic analysis of social interactions deconstructs the evolved loss of schooling behavior in cavefish. PLoS One. 2022;17(4). 10.1371/journal.pone.0265894

100. Yoshizawa M, Robinson BG, Duboué ER, et al. Distinct genetic architecture underlies the emergence of sleep loss and prey-seeking behavior in the Mexican cavefish. BMC Biol. 2015;13:15. 10.1186/s12915-015-0119-3

101. Krause J. The influence of food competition and predation risk on size-assortative shoaling in juvenile chub (*Leuciscus cephalus*). Ethology. 1994;96(2):105–16. 10.1111/j.1439-0310.1994.tb00886.x

102. Pitcher TJ. Functions of shoaling behaviour in teleosts. In: Pitcher TJ, editor. The behaviour of teleost fishes. Springer; 1986. p. 294–337. 10.1007/978-1-4684-8261-4_12

103. Fricke D. Reaction to alarm substance in cave populations of *Astyanax fasciatus* (Characidae, Pisces). Ethology. 1987;76(4):305–8. 10.1111/j.1439-0310.1987.tb00696.x

104. Rétaux S, Elipot Y. Feed or fight: A behavioral shift in blind cavefish. Commun Integr Biol. 2013;6(2). 10.4161/cib.23166

105. Parzefall J, Hausberg C. Ontogeny of the aggressive behaviour in epigean and hypogean populations of *Astyanax fasciatus* (Characidae, Teleostei) and their hybrids. Mem Biospeol. 2001;28:157–61.

106. Espinasa L, Rivas-Manzano P, Pérez HE. A new blind cave fish population of genus *Astyanax*: Geography, morphology, and behavior. Environ Biol Fishes. 2001;62(1):339–44. 10.1023/A:1011852603162

107. Iwata M. Downstream migratory behavior of salmonids and its relationship with cortisol and thyroid hormones: A review. Aquaculture. 1995;135(1-3):131–9. 10.1016/0044-8486(95)01000-9

108. Oliveira RF, Gonçalves DM. Hormones and social behaviour of teleost fish. In: Fish behaviour. 1st ed. CRC Press; 2008. p. 90.

109. Goodson JL, Bass AH. Social behavior functions and related anatomical characteristics of vasotocin/vasopressin systems in vertebrates. Brain Res Rev. 2001;35(3):246–65. 10.1016/s0165-0173(01)00043-1

110. Pierre C, Pradère N, Froc C, et al. A mutation in monoamine oxidase (MAO) affects the evolution of stress behavior in the blind cavefish *Astyanax mexicanus*. J Exp Biol. 2020;223. 10.1242/jeb.226092

111. Moore FL, Boyd SK, Kelley DB. Historical perspective: Hormonal regulation of behaviors in amphibians. Horm Behav. 2005;48(4):373–83. 10.1016/j.yhbeh.2005.05.011

112. Godwin J, Thompson R. Nonapeptides and social behavior in fishes. Horm Behav. 2012;61(3):230–8. 10.1016/j.yhbeh.2011.12.016

113. Thompson RR, Walton JC. Peptide effects on social behavior: Effects of vasotocin and isotocin on social approach behavior in male goldfish (*Carassius auratus*). Behav Neurosci. 2004;118(3):620–6. 10.1037/0735-7044.118.3.620

114. Cabrera-Álvarez MJ. Neural mechanisms of social behaviour and social information use in guppies (Poecilia reticulata) [Doctoral dissertation]. McGill University; 2018.

115. Ataei Mehr B. Effects of the brain nonapeptides arginine-vasotocin and isotocin on shoaling behaviour in the guppy (Poecilia reticulata) [Master’s thesis]. Western University; 2022. https://ir.lib.uwo.ca/etd/8905

116. Lindeyer CM, Langen EMA, Swaney WT, Reader SM. Nonapeptide influences on social behaviour: Effects of vasotocin and isotocin on shoaling and interaction in zebrafish. Behaviour. 2015;152(7-8):897–915. 10.1163/1568539X-00003261

117. Filby AL, Paull GC, Hickmore TF, Tyler CR. Unravelling the neurophysiological basis of aggression in a fish model. BMC Genomics. 2010;11:498. 10.1186/1471-2164-11-498

118. Lema SC, Nevitt GA. Exogenous vasotocin alters aggression during agonistic exchanges in male Amargosa River pupfish (*Cyprinodon nevadensis amargosae*). Horm Behav. 2004;46(5):628–37. 10.1016/j.yhbeh.2004.07.003

119. Semsar K, Kandel FLM, Godwin J. Manipulations of the AVT system shift social status and related courtship and aggressive behavior in the bluehead wrasse. Horm Behav. 2001;40(1):21–31. 10.1006/hbeh.2001.1663

120. Bastian J, Schniederjan S, Nguyenkim J. Arginine vasotocin modulates a sexually dimorphic communication behavior in the weakly electric fish *Apteronotus leptorhynchus*. J Exp Biol. 2001;204(Pt 11):1909–23. 10.1242/jeb.204.11.1909

121. Santangelo N, Bass AH. New insights into neuropeptide modulation of aggression: Field studies of arginine vasotocin in a territorial tropical damselfish. Proc Biol Sci. 2006;273(1605):3085–92. 10.1098/rspb.2006.3683

122. Reddon AR, Voisin MR, O’Connor CM, Balshine S. Isotocin and sociality in the cooperatively breeding cichlid fish, *Neolamprologus pulcher*. Behaviour. 2014;151(10):1389–411. 10.1163/1568539X-00003190

123. Reddon AR, O’Connor CM, Marsh-Rollo SE, Balshine S. Effects of isotocin on social responses in a cooperatively breeding fish. Anim Behav. 2012;84(4):753–60. 10.1016/j.anbehav.2012.07.021

124. O’Connor CM, Marsh-Rollo SE, Aubin-Horth N, Balshine S. Species-specific patterns of nonapeptide brain gene expression relative to pair-bonding behavior in grouping and non-grouping cichlids. Horm Behav. 2016;80:30–8. 10.1016/j.yhbeh.2015.10.015

125. Oldfield RG, Hofmann HA. Neuropeptide regulation of social behavior in a monogamous cichlid fish. Physiol Behav. 2011;102(3-4):296–303. 10.1016/j.physbeh.2010.11.022

126. Greenwood AK, Wark AR, Fernald RD, Hofmann HA. Expression of arginine vasotocin in distinct preoptic regions is associated with dominant and subordinate behavior in an African cichlid fish. Proc Biol Sci. 2008;275(1653):2393–402. 10.1098/rspb.2008.0622

127. Reddon AR, Aubin-Horth N, Reader SM. Wild guppies from populations exposed to higher predation risk exhibit greater vasotocin brain gene expression. J Zool. 2021. 10.1111/jzo.12937

128. Ramsey ME, Fry D, Cummings ME. Isotocin increases female avoidance of males in a coercive mating system: Assessing the social salience hypothesis of oxytocin in a fish species. Horm Behav. 2019;111:112–9. 10.1016/j.yhbeh.2019.03.001

129. Shamay-Tsoory SG, Abu-Akel A. The social salience hypothesis of oxytocin. Biol Psychiatry. 2016;79(3):194–202. 10.1016/j.biopsych.2015.07.020

130. Striepens N, Scheele D, Kendrick KM, Hurlemann R. Oxytocin facilitates protective responses to aversive social stimuli in males. Proc Natl Acad Sci USA. 2012;109(44):18144–9. 10.1073/pnas.1208852109

131. Burkhart JC, Gupta S, Borrego N, Heilbronner SR, Packer C. Oxytocin promotes social proximity and decreases vigilance in groups of African lions. iScience. 2022;25(4):104049. 10.1016/j.isci.2022.104049

132. Smith AS, Agmo A, Birnie AK, French JA. Manipulation of the oxytocin system alters social behavior and attraction in pair-bonding primates, *Callithrix penicillata*. Horm Behav. 2010;57(2):255–62. 10.1016/j.yhbeh.2009.12.004

133. Simpson EA, Sclafani V, Paukner A, et al. Inhaled oxytocin increases positive social behaviors in newborn macaques. Proc Natl Acad Sci USA. 2014;111(19):6922–7. 10.1073/pnas.1402471111

134. Carter GG, Wilkinson GS. Intranasal oxytocin increases social grooming and food sharing in the common vampire bat (*Desmodus rotundus*). Horm Behav. 2015;75:150–3. 10.1016/j.yhbeh.2015.10.006

135. Mooney SJ, Douglas NR, Holmes MM. Peripheral administration of oxytocin increases social affiliation in the naked mole-rat (*Heterocephalus glaber*). Horm Behav. 2014;65(4):380–5. 10.1016/j.yhbeh.2014.02.003

136. Madden JR, Clutton-Brock TH. Experimental peripheral administration of oxytocin elevates a suite of cooperative behaviours in a wild social mammal. Proc Biol Sci. 2010;278(1709):1189–94. 10.1098/rspb.2010.1675

137. Kosfeld M, Heinrichs M, Zak P, Fi schbacher U, Fehr E. Oxytocin increases trust in humans. Nature. 2005;435:673–6. 10.1038/nature0370

138. Bartz JA, Zaki J, Bolger N, Ochsner KN. Social effects of oxytocin in humans: Context and person matter. Trends Cogn Sci. 2011;15(7):301–9. 10.1016/j.tics.2011.05.002

139. Crockford C, Deschner T, Ziegler TE, Wittig RM. Endogenous peripheral oxytocin measures can give insight into the dynamics of social relationships: A review. In: Keebaugh A, Adnari E, Parr LA, editors. Oxytocin’s routes in social behavior: Into the 21st century. Lausanne: Frontiers Media; 2015. p. 40–53.

140. Kelly AM, Goodson JL. Social functions of individual vasopressin–oxytocin cell groups in vertebrates: What do we really know? Front Neuroendocrinol. 2014;35:512–29. 10.1016/j.yfrne.2014.04.005

141. Goodson JL, Thompson RR. Nonapeptide mechanisms of social cognition, behavior, and species-specific social systems. Curr Opin Neurobiol. 2010;20(6):784–94. 10.1016/j.conb.2010.08.020

142. Carneiro L, Oliveira R, Canário A. The effect of arginine vasotocin on courtship behaviour in a blenniid fish with alternative reproductive tactics. Fish Physiol Biochem. 2003;28:241–3. 10.1023/B:FISH.0000030542.31395.8a

143. Pérez-Escudero A, Vicente-Page J, Hinz R, de Polavieja GG. idTracker: Tracking individuals in a group by automatic identification of unmarked animals. Nat Methods. 2014;11(7):743–8. 10.1038/nmeth.2994

144. R Core Team. R: A language and environment for statistical computing. R Foundation for Statistical Computing; 2023. https://www.R-project.org/

145. Miller N, Gerlai R. Automated tracking of zebrafish shoals and the analysis of shoaling behavior. In: Kalueff A, Stewart A, editors. Zebrafish protocols for neurobehavioral research. Humana Press; 2012. Neuromethods, vol 66. 10.1007/978-1-61779-597-8_16

146. Miller N, Gerlai R. From schooling to shoaling: Patterns of collective motion in zebrafish (*Danio rerio*). PLoS One. 2012;7(11). 10.1371/journal.pone.0048865

147. Iwashita M, Yoshizawa M. Social-like responses are inducible in asocial Mexican cavefish despite the exhibition of strong repetitive behavior. eLife. 2021;10. 10.7554/eLife.72463

148. McHenry MJ, Strother JA, van Netten SM. Mechanical filtering by the boundary layer and fluid–structure interaction in the superficial neuromast of the fish lateral line system. J Comp Physiol A. 2008;194(9):795–810. 10.1007/s00359-008-0350-2

149. Reddon AR, O’Connor CM, Nesjan E, et al. Isotocin neuronal phenotypes differ among social systems in cichlid fishes. R Soc Open Sci. 2017;4(5):170350. 10.1098/rsos.170350

150. Guerra AS, Kao AB, McCauley DJ, Berdahl AM. Fisheries-induced selection against schooling behaviour in marine fishes. Proc Biol Sci. 2020;287(1935):20201752. 10.1098/rspb.2020.1752

