## Supplementary Information for "Mechanisms of Social Behavior in the Anti-Social Blind Cavefish (*Astyanax mexicanus*)"

**Table S1**. Means of all measures in all experiments. The table gives the mean measure (IID or NND, in cm; Polarization in unitless values) for each condition in each experiment. The table is divided by experiment (1-4).

Experiment 1 (comparison of zebrafish, sighted AM, blind AM, and theoretical null model):

| | Theory null | Blind AM | Sighted AM | Zebrafish |
|---|---|---|---|---|
| NND (cm) | 10.61 ± 6.40 | 12.80 ± 9.39 | 3.48 ± 2.45 | 4.96 ± 4.69 |
| IID (cm) | 19.34 ± 6.53 | 26.42 ± 7.62 | 6.57 ± 3.31 | 10.06 ± 6.41 |
| Polarization | 0.40 ± 0.20 | 0.40 ± 0.20 | 0.72 ± 0.24 | 0.66 ± 0.25 |

Experiment 2 (comparison of fed and hungry blind AM [compared in the text to control data from Experiment 1]):

| | Hungry | Fed |
|---|---|---|
| NND (cm) | 14.50 ± 8.79 | 14.24 ± 8.81 |
| IID (cm) | 27.85 ± 7.84 | 27.66 ± 7.80 |
| Polarization | 0.39 ± 0.20 | 0.43 ± 0.21 |

Experiment 3 (comparison of drug effects on blind and sighted AM):

| **Blind AM** | Saline | IT+ | IT- | AVT+ | AVT- |
|---|---|---|---|---|---|
| NND (cm) | 13.56 ± 3.92 | 13.75 ± 3.95 | 12.75 ± 3.94 | 12.67 ± 3.70 | 13.34 ± 4.03 |
| IID (cm) | 26.67 ± 4.92 | 27.01 ± 4.53 | 24.84 ± 4.79 | 24.72 ± 4.97 | 26.14 ± 5.24 |
| Polarization | 0.41 ± 0.19 | 0.41 ± 0.19 | 0.41 ± 0.20 | 0.40 ± 0.20 | 0.45 ± 0.20 |
| **Sighted AM** | | | | | |
| NND (cm) | 3.79 ± 1.73 | 3.50 ± 1.53 | 3.94 ± 1.69 | 4.59 ± 1.78 | 3.31 ± 1.19 |
| IID (cm) | 7.24 ± 3.50 | 6.95 ± 2.98 | 7.69 ± 3.71 | 9.53 ± 4.50 | 6.23 ± 2.44 |
| Polarization | 0.74 ± 0.25 | 0.55 ± 0.25 | 0.71 ± 0.24 | 0.68 ± 0.26 | 0.71 ± 0.25 |

[continued on next page]

Experiment 4 (comparison of higher doses of AVT+ and IT- on schooling in blind AM [compared in the text to saline and 10 µg/g data from Experiment 3]):

| | AVT+ @ 20 µg/g | AVT+ @ 40 µg/g | IT- @ 40 µg/g |
|---|---|---|---|
| NND (cm) | 13.59 ± 7.79 | 14.64 ± 9.05 | 12.87 ± 7.57 |
| IID (cm) | 25.91 ± 6.96 | 28.45 ± 7.66 | 24.21 ± 7.00 |
| Polarization | 0.39 ± 0.20 | 0.38 ± 0.20 | 0.40 ± 0.20 |

**Table S2**: ANOVA Post-hoc test results for Experiment 1. Post-hoc tests were run using the Bonferroni correction. Each cell gives the T statistic for the test along with a p-value and the value of Cohen's D. Each sub-table gives results for a different measure (IID, NND, or Polarization).

| **IID** | Blind AM | Sighted AM |
|---|---|---|
| Sighted AM | T(124.1) = -53.1, p < 0.00001, D = 9.11 | |
| Zebrafish | T(138) = -35.2, p < 0.00001, D = 5.96 | T(114.7) = 8.35, p < 0.00001, D =1.42 |
| **NND** | | |
| Sighted AM | T(113.4) = -41.7, p < 0.00001, D = 7.06 | |
| Zebrafish | T(138) = -31.0, p < 0.00001, D = 5.24 | T(120.4) = 7.26, p < 0.00001, D = 1.24 |
| **Polarization** | | |
| Sighted AM | T(96.0) = -20.4, p < 0.00001, D = 3.70 | |
| Zebrafish | T(118.5) = -18.4, p < 0.00001, D = 3.10 | T(128) = -3.00, p = 0.004, D = 0.53 |

**Table S3**: ANOVA Post-hoc test results for Experiment 2. Post-hoc tests were run using the Bonferroni correction. Control data are taken from Experiment 1. Each cell gives the T statistic for the test along with a p-value and the value of Cohen's D. Each sub-table gives results for a different measure (IID, NND, or Polarization).

| **IID** | Hungry | Fed |
|---|---|---|
| Control | T(128) = -4.06, p = 0.00008, D = 0.71 | T(118) = -3.57, p = 0.0005. D = 0.66 |
| Fed | T(108) = -0.41, p = 0.69, D = 0.08 | |
| **NND** | | |
| Control | T(128) = -6.92, p < 0.00001, D = 1.22 | T(118) = -5.76, p < 0.00001, D = 1.07 |
| Fed | T(108) = -0.95, p = 0.34, D = 0.18 | |
| **Polarization** | | |
| Control | T(128) = 1.45, p = 0.15, D = 0.25 | T(118) = -2.35, p = 0.02, D = 0.43 |
| Fed | T(108) = -4.05, p = 0.0001, D = 0.77 | |

**Table S4**: ANOVA Post-hoc test results for Experiment 3, for comparisons between strains (blind and sighted). Post-hoc tests were run using the Bonferroni correction. Each cell gives the T statistic for the test along with a p-value and the value of Cohen's D. Each row gives results for a different measure (IID, NND, or Polarization [POL]).

| | Saline | IT+ | IT- | AVT+ | AVT- |
|---|---|---|---|---|---|
| **IID** | T(78) = 42.0, p < 0.00001, D = 9.71 | T(68) = 47.47, P < 0.00001, D = 11.46 | T(68) = 29.24, P < 0.00001, D = 7.06 | T(68) = 23.83, P < 0.00001, D = 5.75 | T(45.4) = 44.59, P < 0.00001, D = 11.48 |
| **NND** | T(78) = 42.3, P < 0.00001, D = 9.77 | T(68) = 42.00, P < 0.00001, D = 10.12 | T(42.2) = 23.76, p < 0.00001, D = 6.20 | T(67.3) = 34.77, P < 0.00001, D = 7.95 | T(38.5) = 36.54, P < 0.00001, D = 9.69 |
| **POL** | T(34.2) = 13.34, P < 0.00001, D = 3.72 | T(61.3) = 6.52, P < 0.00001, D = 1.45 | T(51.6) = 13.57, P < 0.00001, D = 2.93 | T(34.7) = 11.18, P < 0.00001, D = 3.02 | T(63.7) = 11.18, P < 0.00001, D = 2.50 |

**Table S5**: ANOVA Post-hoc test results for Experiment 3, for comparisons between saline and the drug treatment conditions. Post-hoc tests were run using the Bonferroni correction. Each cell gives the T statistic for the test along with a p-value and the value of Cohen's D. Each row gives results for a different measure (IID, NND, or Polarization). The top part of the table gives results for blind AM, the bottom for sighted AM. Each drug condition is being compared to the saline condition.

| **Blind** | IT+ | IT- | AVT+ | AVT- |
|---|---|---|---|---|
| **IID** | T(78) = -0.75,<br>p = 0.45,<br>D = 0.17 | T(78) = 3.70,<br>p = 0.0004,<br>D = 0.85 | T(88) = 4.35,<br>p = 0.00004,<br>D = 0.92 | T(78) = 1.09,<br>p = 0.28,<br>D = 0.25 |
| **NND** | T(78) = -0.80,<br>P = 0.42,<br>D = 0.19 | T(40.3) = 2.17,<br>P = 0.04,<br>D = 0.57 | T(88) = 3.79,<br>P = 0.0003,<br>D = 0.80 | T(78) = 0.72,<br>P = 0.47,<br>D = 0.17 |
| **Polarization** | T(78) = 0.16,<br>P = 0.87,<br>D = 0.04 | T(78) = -0.07,<br>P = 0.94,<br>D = 0.02 | T(88) = 0.85,<br>P = 0.40,<br>D = 0.18 | T(46.7) = -2.72,<br>P = 0.009,<br>D = 0.68 |
| **Sighted** | | | | |
| **IID** | T(68) = 0.64,<br>p = 0.52,<br>D = 0.15 | T68) = -0.79,<br>p = 0.43,<br>D = 0.19 | T(58) = -3.37,<br>p = 0.001,<br>D = 0.87 | T(68) = 2.41,<br>p = 0.02,<br>D = 0.58 |
| **NND** | T(61.9) = 1.23,<br>P = 0.22,<br>D = 0.30 | T(68) = -0.67,<br>P = 0.50,<br>D = 0.16 | T(58) = -3.64,<br>P = 0.0006,<br>D = 0.94 | T(68) = 2.46,<br>P = 0.02,<br>D = 0.59 |
| **Polarization** | T(68) = 6.51,<br>P < 0.00001,<br>D = 1.57 | T(68) = 0.76,<br>P = 0.45,<br>D = 0.18 | T(58) = 1.81,<br>P = 0.07,<br>D = 0.47 | T(68) = 0.90,<br>P = 0.37,<br>D = 0.22 |

**Table S6**. ANOVA Post-hoc test results for Experiment 4, comparing each drug concentration to saline (saline results taken from Experiment 3). Post-hoc tests were run using the Bonferroni correction. Each cell gives the T statistic for the test along with a p-value and the value of Cohen's D. Each row gives results for a different measure (IID, NND, or Polarization).

| | AVT+ @ 20 μg/g | AVT+ @ 40 μg/g | IT- @ 40 μg/g |
|---|---|---|---|
| **IID** | T(78) = 1.26, P = 0.21, D = 0.29 | T(45.8) = -3.12, P = 0.003, D = 0.79 | T(86) = 5.06, P < 0.00001, D = 1.01 |
| **NND** | T(78) = -0.36, P = 0.72, D = 0.08 | T(78) = -4.10, P = 0.0001, D = 0.95 | T(87.3) = 2.62, P = 0.01, D = 0.52 |
| **Polarization** | T(78) = 2.47, P = 0.02, D = 0.57 | T(78) = 3.03, P = 0.003, D = 0.70 | T(98) = 1.53, P = 0.13, D = 0.31 |

**Figure S1**. Frame of video from a trial of blind AM in Experiment 1. The tank was 60 cm in diameter.

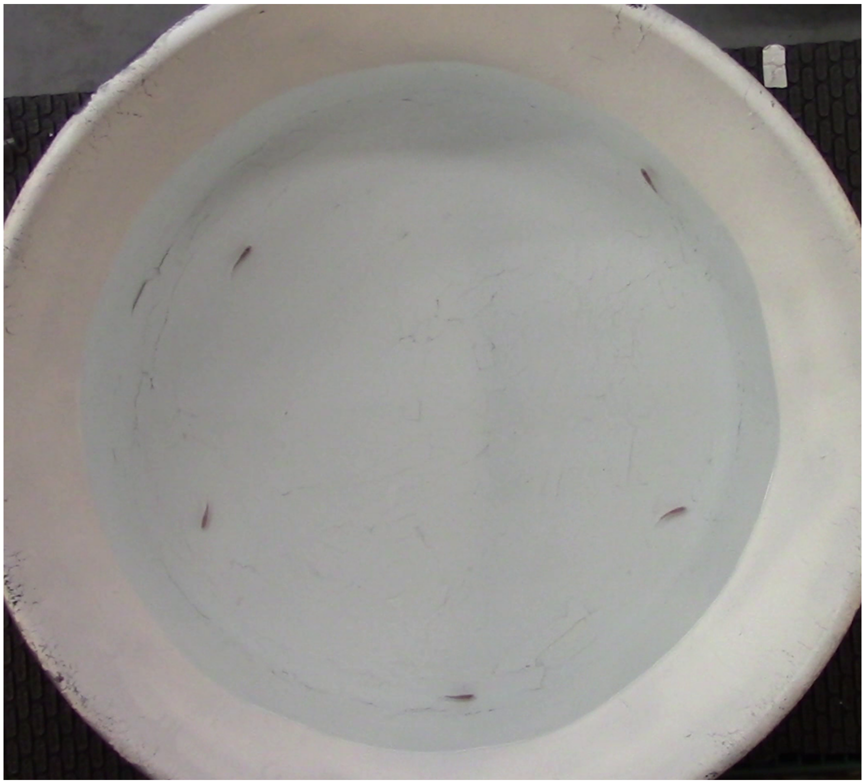

**Video S1**: sample trial of blind AM from Experiment 1.

**Video S2**: sample trial of sighted AM from Experiment 1.
